# The multifurcating skyline plot

**DOI:** 10.1101/356097

**Authors:** Patrick Hoscheit, Oliver G. Pybus

**Affiliations:** MaIAGE, INRA, Universit Paris-Saclay, 78350 Jouy-en-Josas, France; Department of Zoology, University of Oxford, Oxford OX1 3PS, UK

## Abstract

A variety of methods based on coalescent theory have been developed to infer demographic history from gene sequences sampled from natural populations. The “skyline plot” and related approaches are commonly employed as flexible prior distributions for phylogenetic trees in the Bayesian analysis of pathogen gene sequences. In this work we extend the classic and generalised skyline plot methods to phylogenies that contain one or more multifurcations (*i.e.* hard polytomies). We use the theory of Λ-coalescents (specifically, Beta(2*α*,*α*)-coalescents) to develop the “multifurcating skyline plot”, which estimates a piecewise constant function of effective population size through time, conditional on a time-scaled multifurcating phylogeny. We implement a smoothing procedure and extend the method to serially-sampled (heterochronous) data, but we do not address here the problem of estimating trees with multifurcations from gene sequence alignments. We validate our estimator on simulated data using maximum likelihood and find that parameters of the Beta(2*α*,*α*)-coalescent process can be estimated accurately. Lastly we apply the multifurcating skyline plot to a molecular clock phylogeny of 1,610 Ebola virus sequences from the 2014-2016 West African outbreak. We artificially collapse short branches in this empirical phylogeny in order to mimic different levels of multifurcation and show that variance in the reproductive success of the pathogen through time can be estimated by combining the skyline plot with epidemiological case count data.

## Introduction

The field of *phylodynamics* is concerned with the study of how processes acting at the population level shape the genetic diversity of of gene or genome sequences sampled from natural populations. Phylodynamic methods are frequently applied to pathogen populations and used to test hypotheses concerning the epidemiology and transmission of infectious disease [e.g. (Pybus and Rambaut, 2009)]. Viruses in particular have been the subject of great attention, since their high mutation rates rapidly generate genetic diversity, even on short time scales, and because increasingly large numbers of virus genome sequences from viral epidemics are avilable for analysis [e.g. (Biek *et al.*, 2015)]. Phylodynamic approaches are also used in other fields, such as macroevolution, anthropology, and ancient DNA research [e.g. (Drummond *et al.*, 2003)], in order to understand how dynamical processes gave rise to the patterns of ancestry and diversity observed in biological systems.

Phylodynamic analysis of sampled gene sequences relies crucially on “tree-generating models”, which describe how phylogenies, genealogies or trees (we use these terms interchangeably) are related to the population dynamic processes that generated them. Among these models, coalescent approaches are widely used because they provide a mathematically simple framework to relate the demography of a viral population to its sample genealogy. Mathematically, coalescent theory relies on asymptotic properties of Wright-Fisher reproduction models that represent large, constant-sized populations. The distribution of sample genealogies from such populations are described by the so-called Kingman coalescent process (Kingman, 1982). The Kingman coalescent has been shown to describe the genealogy of many models in population genetics and has been extended to incorporate a number of biological processes, including population size change (Griffiths and Tavare, 1994), selection (Kaplan *et al.*, 1988), recombination (Hudson and Kaplan, 1988), longitudinal sampling (Rodrigo *et al.*, 1999) and population structure (Takahata, 1988).

The Kingman coalescent has been used to develop a variety of statistical methods that aim to infer the history of population size from an observed sample phylogeny. One such approach that is commonly used is the “skyline plot” and related methods (Ho and Shapiro, 2011), which model population size change as a piecewise constant function through time. In this paper, we extend and generalise the skyline plot family of methods beyond the Kingman coalescent.

The standard Kingman coalescent (Kingman, 1982) describes the genealogy of *n* individuals sampled at random from a population of size *N* ≫ *n*, using a bifurcating ultrametric tree *T*_*n*_ with *n* leaves (tree tips). In this paper, we will consider genealogies obtained by sampling from large populations whose sizes vary over time. In contrast to standard Kingman coalescent theory, we will consider populations with high variance in the number of offspring per individual (sometimes called the offspring distribution) which may lead to multifurcations (i.e. nodes with degree > 3) in the sample genealogy that can no longer be ignored. To achieve this we consider a more general class of models called Λ-coalescent processes, a family of random trees discovered by Sagitov (Sagitov, 1999) and fully described by Pitman (Pitman, 1999). In contrast to the Kingman coalescent, under which trees are binary and strictly bifurcating, Λ-coalescent trees can contain multifurcations.

Here we show how to calculate the likelihood of multifurcating genealogies under Λ-coalescent models, given a function describing effective population size. We derive a new estimate of effective population size, which we call the multifurcating skyline plot, and extend this to longitudinal (serial) sampling. Using simulated data, we show that, in the case of Beta (2 − *α*, *α*)-coalescents, we can estimate the key *α* parameter that describes the propensity of the sample genealogy to contain multifurcations (*α* = 2 corresponds to the Kingman coalescent, whilst *α* = 1 represents the Bolthausen-Sznitman coalescent). We also show that effective population sizes also can be estimated accurately. Note that this study aims to develop methods that estimate effective population size from a single, prespecified multifurcating genealogy. We leave for future work the problem of statistically inferring multifurcating trees from an empirical gene sequence alignment.

We apply our methods to a phylogeny estimated from virus genomes sampled and sequenced during the 2014-2015 Ebola virus epidemic in West Africa (Dudas *et al.*, 2017). Using a heuristic approach, we aggregate nodes in this empirical genealogy in order to generate a pseudo-multifurcating tree, which was analyzed subsequently using the multifurcating skyline plot. The plot leads to a different estimate of the effective population size of the Ebola virus epidemic than that obtained using Kingman coalescent-based methods. We further use our method to infer temporal fluctuations in the variance of the offspring distribution of the epidemic’s infection process.

## Λ-COALESCENTS

### A. Mathematical properties

Coalescents are random trees that describe the genealogy of a small number of individuals sampled from a larger population. The class of Λ-coalescents is parametrised by a probability measure Λ(*dx*) on the interval [0; 1]. In most cases, these probability measures will be absolutely continuous with respect to the Lebesgue measure, *i.e*. Λ(*dx*) = λ(*x*)*dx*, for some nonnegative function λ(*x*) such that 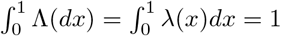.

Under the Kingman coalescent, sample trees are almost surely binary, However, trees under the Λ-coalescent are in general multifurcating (i.e. contain at least one node with degree larger than 3). The distribution of such trees can be described as follows: start with *n* sample lineages at time 0. Note that time is represented backwards hence time 0 represents the present (or, more specifically, the time of the most recent tree tip). Consider, for each *k*-tuple of lineages with 2 ≤ *k* ≤ *n*, an independent exponential random variable with parameter

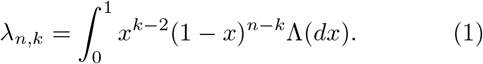

Then, if *T*_*n*_ is the minimum of these random varia Pbles, *T*_*n*_ is exponentially distributed with parameter 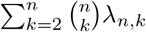. At time *T*_*n*_, choose the number *K* of coalescing lineages with probability

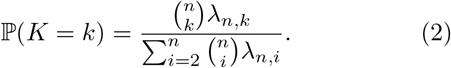

Finally, choose *K* lineages at random from the set of n extant lineages, and collapse them into a single lineage. Continue this process with the remaining *n* − *K* + 1 lineages until only one lineage is left. In other words, each *k*-tuple of lineages coalesces at rate λ_*n*, *k*_, independently of all other tuples. Note that time is represented *backwards* since time 0 is the present. To better understand the role played by the Λ measure in the distribution of multifurcations, equation (2) can also be interpreted in the following way: assume that 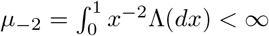, so that *F*_1_ (*dx*) = *x*^−2^Λ(*dx*)/*μ*−2 is a probability measure. Then, at coalescence time *T*_*n*_, let *p* ∈ (0,1] be sampled according to *F*_1_; select coalescing lineages from the *n* extant lineages independently with probability *p*. For more details on this construction (sometimes dubbed the *paintbox* construction by analogy with the construction used by Kingman) see (Pitman, 1999).

In equation (1), it is easy to see that when Λ = *δ*_0_, the Dirac mass at 0, then λ_*n*,*k*_ = 0 for 3 ≤ *k* ≤ *n*, and λ_*n*,2_ = 1. Thus, if the probability measure Λ is concentrated solely at 0, then the Λ-coalescent process is strictly binary and identical to the Kingman coalescent. However, the paintbox construction scheme does not apply to this case, since 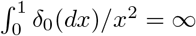.

### B. Variable population size

For a given set of *n* individuals sampled from a large populations, we can define the *coalescent effective population size*, *N*_*e*_, which is the size of an ideal population exhibiting the same amount of genetic diversity under the Wright-Fisher reproduction model as the population under study. Indeed, one can recover a Kingman coalescent from the genealogy of *n* individuals in a Wright-Fisher model of size *N* by rescaling time by a factor of *N*_*e*_ = *N* (Wakeley, 2009). For more details on the different notions of effective population size see (Sjödin *et al.*, 2005).

For Λ-coalescents, the analog of the Wright-Fisher model is the *Cannings model* (Cannings, 1974). In the Cannings model, all *N* individuals in one generation reproduce according to the same offspring distribution (see Fig. 1). If *ν*_1_,…, *ν*_*N*_ are the number of offspring of individuals 1,…, *N* respectively, we need to have *ν*_1_ +…+ *ν*_*N*_ = *N* in order to keep population size constant. A further condition is *exchangeability*, meaning that the distribution of the vector (*ν*_1_,…,*ν*_*N*_) is invariant under permutations. This implies in particular that all individuals have the same offspring distribution (i.e. the same propensity to reproduce). Hence, their common expectation is 𝔼[*ν*_1_] =…= 𝔼[*ν*_*N*_] = 1. Let *σ*^2^(*N*) be their common variance. We can recover the Wright-Fisher model as a specific case of the Cannings model by taking (*ν*_1_,…, *ν*_*N*_) to be the multinomial distributions with parameters (*N*; 1/*N*,…,1/*N*).

**FIG. 1.**
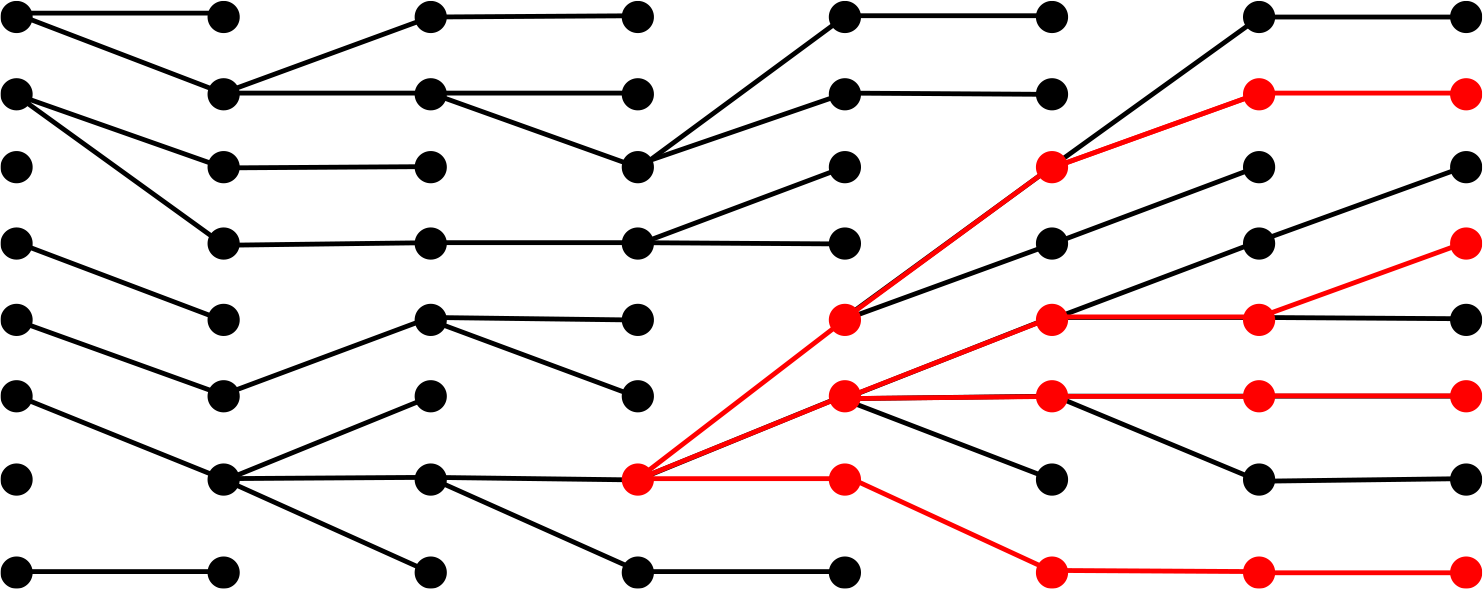
An ancestral genealogy (in red) of *n* = 4 lineages sampled from a Cannings model with constant population size *N* = 8 (black dots). The horizontal axis represents time and the lines indicate ancestry.

Suppose that the population size is large (*N* → ∞). If the variance in offspring number converges to a fixed finite value as *N* increases (*σ*^2^(*N*) → *σ*^2^ < ∞), then the genealogy of *n* lineages randomly sampled from the Cannings model of size N, rescaled by *N*_*e*_ = *N*/σ^2^, still converges to the Kingman coalescent as in the Wright-Fisher case. If we relax this convergence condition, but still keep *σ*^2^(*N*) = *o*(*N*), under some additional technical conditions (see Theorem 3.1 in (Sagitov, 1999)), then the sample genealogy converges to a Λ-coalescent when rescaled by the factor *N*/σ^2^(*N*). By analogy with the Wright-Fisher case, we can call this the Λ-effective population size. Since the Cannings model counts time in units of generations, this means that, when dealing with a real population that exhibits Cannings-like reproduction, it is necessary to count time in units of *N*_*e*_ = *N*/σ^2^(*N*) generations in order to observe Λ-coalescent-like behavious. We can thus define a Λ-coalescent process in continuous time, given an effective population size function (*N*_*e*_(*t*), *t* ≥ 0) by analogy with the variable population size coalescent described in (Griffiths and Tavare, 1994).

Given a tree 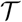 with *n* tips we define 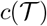 as the number of coalescences, that is, the number of internal nodes of 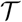 If 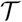 is a binary tree, then 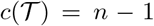. We denote the coalescence times as 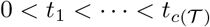. For convenience, we denote present as *t*_0_ = 0. During the interval [*t*_*i*−1_, *t*_*i*_), with 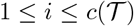, let *n*_*i*_ be the number of extant lineages. By definition, we always have *n*_1_ = *n*. Finally, for each 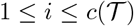, let *k*_*i*_ be the number of lineages involved in the coalescence at time *t*_*i*_ (see Fig. 2 for an example of this notation). Again by definition, we have *n*_*i*+1_ = *n*_*i*_ – *k*_*i*_ + 1.

**FIG. 2.**
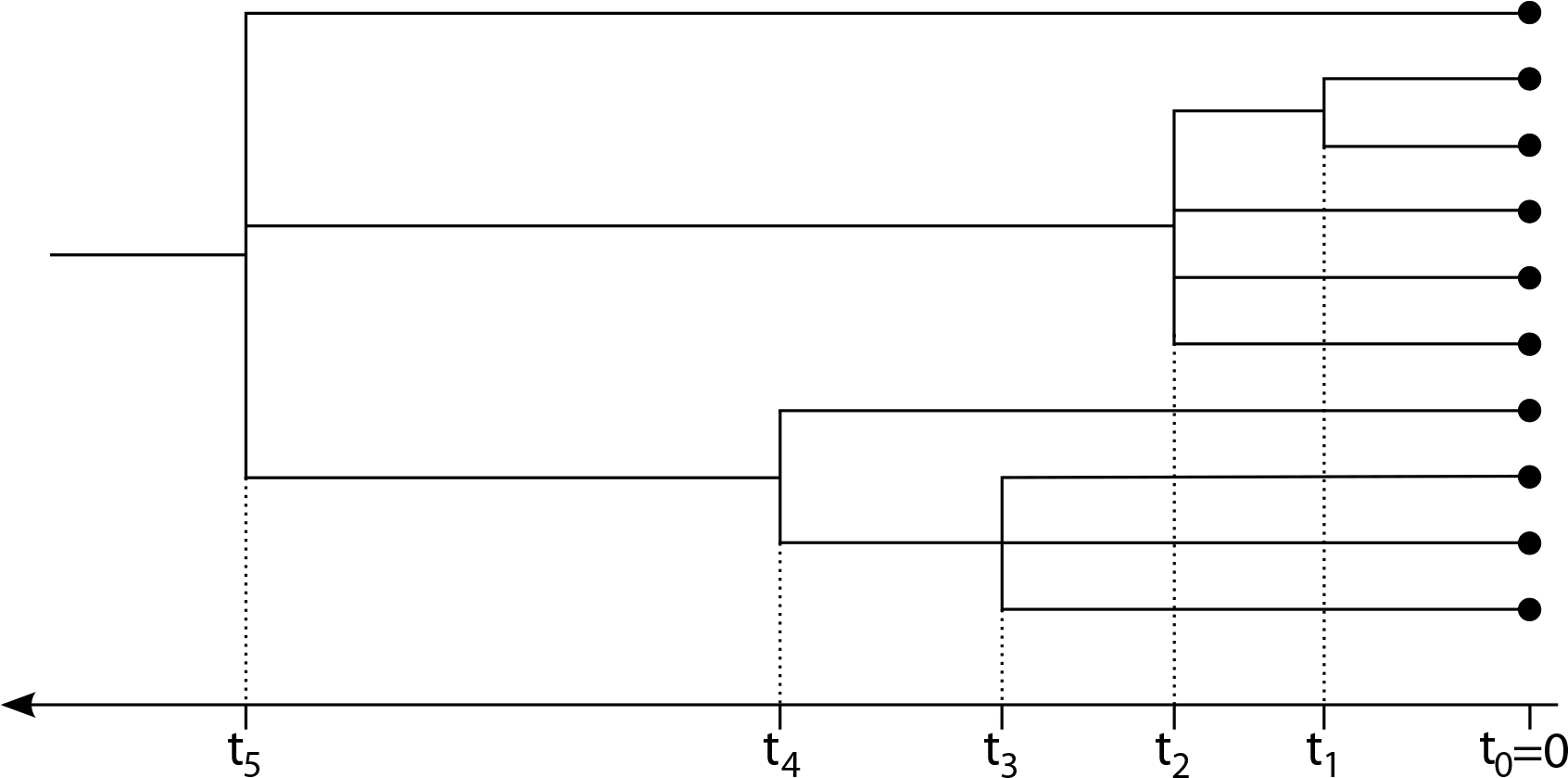
A multifurcating, ultrametric sample tree with *n* =10 tips. In this case, the number of lineages involved in each coalescence event is *k*_1_ = 2, *k*_2_ = 4, *k*_3_ = 3, *k*_4_ = 2, *k*_5_ = 3. The number of extant lineages is *n*_1_ = 10 during [0, *t*_*1*_), *n*_2_ = 9 during [*t*_1_, *t*_2_),…, *n*_5_ = 3 during [*t*_4_,*t*_5_).

The likelihood of a tree under the Λ-coalescent model, given the Λ distribution, is easy to write down thanks to the Markovian description of the process given above. It can in fact be decomposed in product form, since the waiting times between coalescent events are conditionally independent. Given the number of lineages *n*_*i*_ on an intercoalescent interval [*t*_*i*-1_,*t*_*i*_), the waiting time is distributed as an exponential random variable with parameter

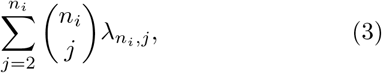

hence the likelihood of observing an interval of length (*t*_*i*_ − *t*_*i*−1_) is

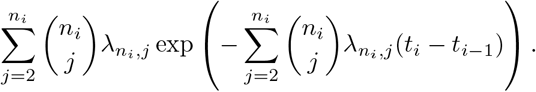

Then, at each coalescent time, the likelihood of seeing exactly *k*_*i*_ of the *n*_*i*_ lineages coalesce is, according to (2),

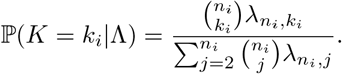

This gives, for the complete likelihood:

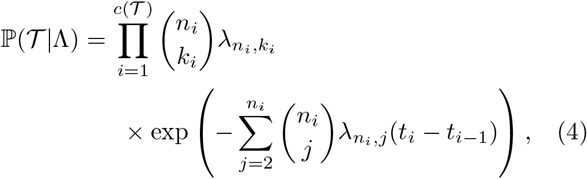

with the λ_*n*,*k*_ depending on Λ as in (1). Note that when taking Λ = *δ*_0_ (i.e. the Kingman coalescent) the λ_*n*,*k*_ are all zero except λ_2,2_ = 1, so that, as expected, we get a zero likelihood for non-binary trees (i.e. trees with at least one *k*_*i*_ > 2) under the Kingman coalescent. Following Griffiths & Tavaré (Griffiths and Tavare, 1994), we can now take into account the effective population size as a time-change of the Λ-coalescent, and write the likelihood of the tree given an effective population size function (*N*_*e*_(*t*),*t* ≥ 0) under a Λ-coalescent model:

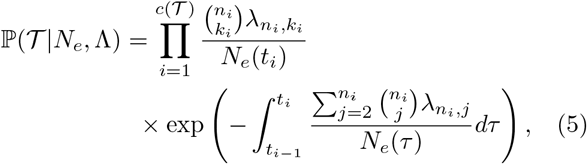

In this formula, the population size function (*N*_*e*_(*t*), *t* ≥ 0) is assumed to be deterministic (see (Kaj and Krone, 2003) for a treatment of Kingman coalescents with stochastically-varying population size). The continuous-time model defined by (5) can be obtained as scaling limit of a Cannings model with variable population size, as shown in (Möhle, 2002). This result, as in the constant-population size case, shows that the effective population size function *N*_*e*_ is related to the parameters of the offspring distribution of a Cannings model through *N*_*e*_(*t*) = *N*(*t*)/σ^2^(*t*), where *N*(*t*) and *σ*^2^ (*t*) are the census population size and the offspring variance, respectively, of the Cannings model at time *t* in the coalescent timescale. This relationship will be useful later in this paper when interpreting estimated effective population sizes. Under thr variable population size model, the change in timescale necessary to transform from generation counts to coalescent time is a little more involved than that for constant population size; details can be found in (Möhle, 2002).

## II. MAXIMUM-LIKELIHOOD ESTIMATION UNDER THE Λ-COALESCENT

Using the likelihood formula (5) and given a time-scaled non-binary tree 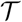, we will now show how to estimate certain features of the process that generated it using maximum likelihood. In the Discussion, we will explore other uses of this likelihood, for instance its use as a tree prior in Bayesian phylogenetic inference frameworks, such as that implemented in BEAST (Drummond and Rambaut, 2007).

### A. Estimation of the Λ measure

We first consider the problem of estimating the Λ parameter directly from an observed genealogy. In the general case the relevant parameter space (i.e. the space of probability measures on [0,1]) is too large to enable direct inference of Λ (see (Koskela *et al.*, 2018) for more advanced techniques). Therefore, we will limit ourselves here to the one-parameter family of `Beta` (2 − *α*, *α*)-coalescents, with *α* ∈ [1, 2]. This corresponds to the case of Λ-coalescents where the Λ measure is the Beta distribution with parameters 2 – *α* and *α*, with *α* ∈ [1, 2):

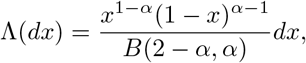

where *B*(*x*,*y*) = Γ(*x*)Γ(*y*)/Γ(*x* + *y*) is the Beta function. By continuity, this can be extended to *α* = 2, in which case the Beta distribution collapses to the Dirac measure at 0, thus recovering the Kingman coalescent. For *α* ∈ [1, 2), the coalescence rates are given by:

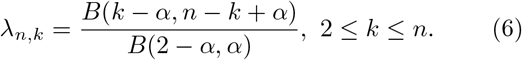

Beta coalescent processes have been extensively studied, starting with Schweinsberg (Schweinsberg, 2003), who described how they can be recovered from the genealogy of certain supercritical branching processes. Similarly, the connection with so-called *α*-stable continuous-state branching processes (Birkner *et al.*, 2005) has enabled the study of many features of the family of Beta-coalescents (Berestycki *et al.*, 2007, 2008; Kersting *et al.*, 2014).

We investigated the problem of inferring the *α* parameter by using trees simulated under the `Beta`(2 − *α*, *α*)-coalescent process with constant effective population size, for three different *α* values, specifically *α* = 1:2; *α* = 1:5; *α* = 1:8. In each case, we simulated 1,000 trees with *n* = 100, 500 and 1000 simultaneously-sampled tips. We then estimated the *α* parameter independently for each tree by maximizing the likelihood (5):

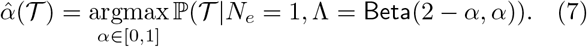

The results of this maximum-likelihood estimation are summarized in Table I and shown in Fig. 3. The procedure yields an effectively unbiased estimator, with decreasing variance as the number of tips increases. Furthermore, for a fixed number of tips, the lowest variance was obtained for true values of *α* close to 2, since the likelihood of trees with non-binary nodes converges to 0 as *α* → 2, so that likelihood landscapes become more peaked. These results are encouraging because they indicate that features of the Λ measure might be identifiable from phylogenies.

**TABLE I.** Empirical mean, bias and variance of maximum-likelihood estimates of the *α* parameter of the Beta-coalescent process.

|  | Number of tips | Mean | Bias | Variance |
| --- | --- | --- | --- | --- |
| $\alpha = 1.2$ | 100 | 1.2004 | $-4 \times 10^{-4}$ | $2.6 \times 10^{-3}$ |
| | 500 | 1.1999 | $-4.8 \times 10^{-5}$ | $2.8 \times 10^{-4}$ |
| | 1000 | 1.1995 | $-4.7 \times 10^{-4}$ | $1.2 \times 10^{-4}$ |
| $\alpha = 1.5$ | 100 | 1.499 | $-1.3 \times 10^{-2}$ | $1.7 \times 10^{-3}$ |
| | 500 | 1.499 | $-1.7 \times 10^{-4}$ | $1.7 \times 10^{-4}$ |
| | 1000 | 1.500 | $1.7 \times 10^{-4}$ | $6.3 \times 10^{-5}$ |
| $\alpha = 1.8$ | 100 | 1.797 | $-2.5 \times 10^{-3}$ | $1.1 \times 10^{-3}$ |
| | 500 | 1.800 | $4.2 \times 10^{-4}$ | $1.1 \times 10^{-4}$ |
| | 1000 | 1.799 | $-2.0 \times 10^{-4}$ | $4.0 \times 10^{-5}$ |

**FIG. 3.**
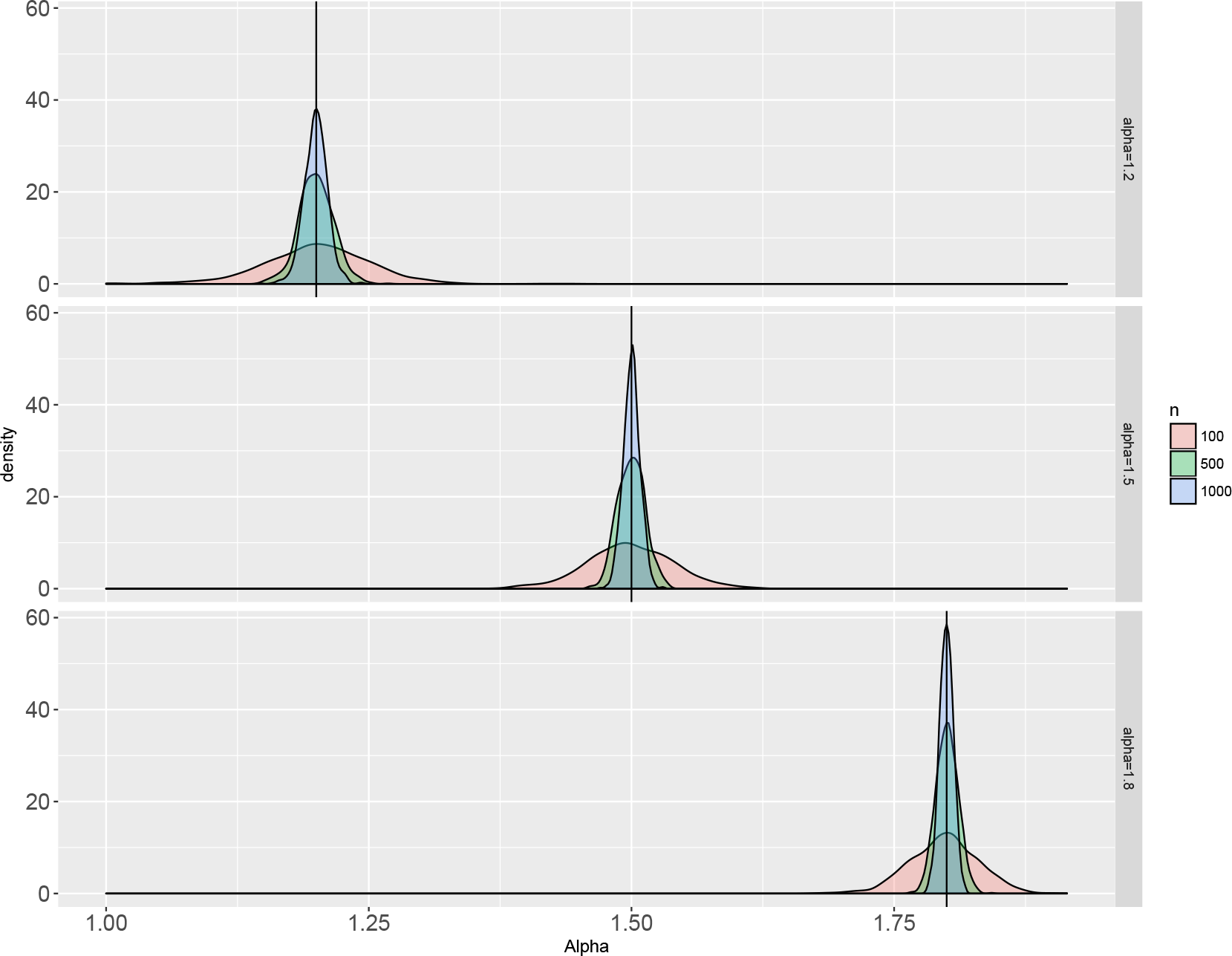
Empirical probability density of maximum-likelihood estimates of *α* ∈ [1; 2] for 1,000 trees simulated under Beta(2 – *α*, *α*)-coalescents, with *α* = 1:2; 1:5 and 1:8, respectively. The color of the density plots corresponds to the number *n* of tips in the simulated trees.

### B. Demographic inference using the skyline plot

We now turn to the problem of estimating ancestral effective population size, using a sample tree as data. Following the classic skyline plot estimator (Pybus *et al.*, 2000), we assume that the duration of inter-coalescent intervals are known without error. We further assume that the degree of each internal node is known precisely. Although both of these variables are uncertain when trees are estimated from empirical data, we leave for future work the problem of incorporating uncertainty in node degree into phylodynamic inference (see Discussion).

It is possible, for a given probability measure Λ, and assuming that population size is constant between coalescent events, to maximize the likelihood function (5). This is made easy by the product form of the likelihood and yields the multifurcating skyline plot estimator:

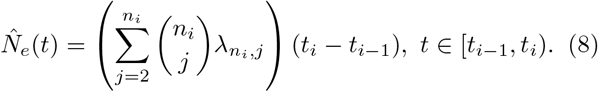

Notice that when Λ = *δ*_0_, the Dirac measure at 0, all the 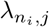 are equal to 0, except for *j* = 2, for which 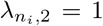, so that we recover the “classic skyline plot” estimate (Pybus *et al.*, 2000) for binary trees:

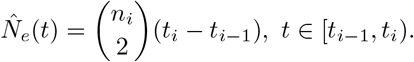

Finally, notice that equation (8) implies that the skyline plot estimator is expressed in the same units as the intercoalescent lengths *t*_*i*_ − *t*_*i*−1_. Hence, if the underlying tree is scaled in real time units, it is necessary to divide the skyline plot estimate by the generation time to recover an estimate in population size units. For a detailed analysis of how varying generation time can affect skyline estimates, see (Volz, 2012).

#### Composite internode intervals

The multifurcating skyline plot (8) is a piecewise constant function on inter-coalescent intervals and there can be a large number of such intervals when the number of tips is large, potentially leading to very noisy estimates. To mitigate this over-fitting, we can use the same interval-merging technique as that employed by the “generalized skyline plot” (Strimmer and Pybus,2001). Given a threshold parameter *ϵ* > 0, we consider all inter-coalescent intervals with length smaller than *ϵ*. We then join those intervals with their neighboring intervals earlier in time (closer to the root), starting with the interval closest to the tips. If the ensuing interval is still of length smaller than *ϵ*, we continue this procedure until there are no more intervals with length smaller than *ϵ*. Note that the degree of internal nodes is unchanged. This yields an interval partition of 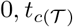. which we denote by 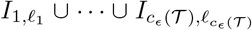 With this notation, 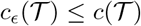 is the number of such composite intervals, and for 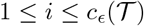, *l*_*i*_ is the number of inter-coalescent intervals joined together to form 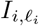.

For each 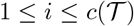, we then independently estimate a single effective population size value for the wholecomposite interval 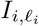, using (5). Due to the product form of the likelihood, it is easy to see that the maximum-likelihood estimator for the composite interval 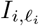 is the mean of the maximum-likelihood estimators (8) for each of the *l*_*i*_ inter-coalescent intervals (note that the original “generalized skyline plot” of (Strimmer and Pybus, 2001) uses a method-of-moments estimator, which leads to a harmonic mean rather than an arithmetic mean).

Thus this composite-interval approach can compute an estimate of effective population size change with fewer parameters. As suggested in (Strimmer and Pybus, 2001), we can then optimize over ϵ ≥ 0 by a model selection statistic such as the corrected Akaike Information Criterion (Hurvich and Tsai, 1989):

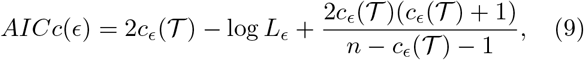

where *L*_*ϵ*_ is the likelihood of the tree given the composite interval skyline estimate with parameter *ϵ*, as computed by (5).

#### Serially-sampled trees

In some biological contexts, such as rapidly evolving pathogens or ancient DNA, sequences are sampled at different times and measurable amounts of sequence change occur between the sampling times (Drummond *et al.*, 2003). As described in (Rodrigo *et al.*, 1999), coalescent estimates can be extended to this framework, by making the assumption that the sampling process is independent of the population dynamics. For the purpose of simplicity, in this paper we also make this independence assumption. We further assume that effective population size does not change at sampling times and that sampling never occurs at coalescence times.

To be precise, let us consider an inter-coalescent interval [*t*_*i*−1_,*t*_*i*_), and assume we have *n*_*i*_ extant lineages at time *t*_*i*−1_. Assume we have sampling times 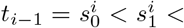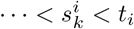, and for 1 ≤ *j* ≤ *k*, let *n*(*j*) be the number of extant lineages on the interval ending with 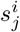 (with *n*(1) = *n*_*i*_). Finally, let *n*(*k* + 1) be the number of lineages on the coalescence interval [*s*_*k*_,*t*_*i*_). Fig. II.B provides a graphical example of a serially-sampled tree and its associated indices.

When computing the likelihood of a serially-sampled tree under a given Λ-coalescent model, given an effective population size function, we need to take into account the intervals of the form 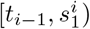 and 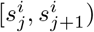 that end with a sampling event. On those intervals thereare no coalescences, so that we need to consider the likelihood of no coalescences occuring during that time. The contribution to the likelihood of the inter-coalescent interval [*t*_*i*-1_,*t*_*i*_) is then:

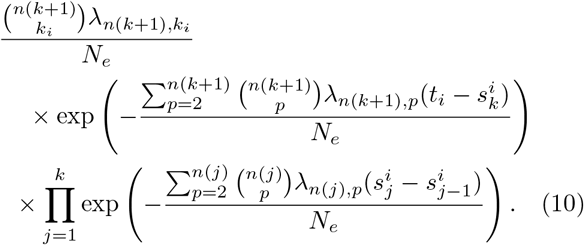

Again, maximizing this in *N*_*e*_ gives the maximum likelihood skyline estimator on the interval [*t*_*i*-1_,*t*_*i*_):

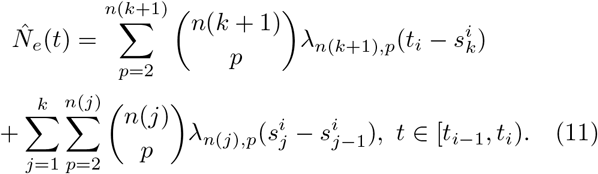

**FIG. 4.**
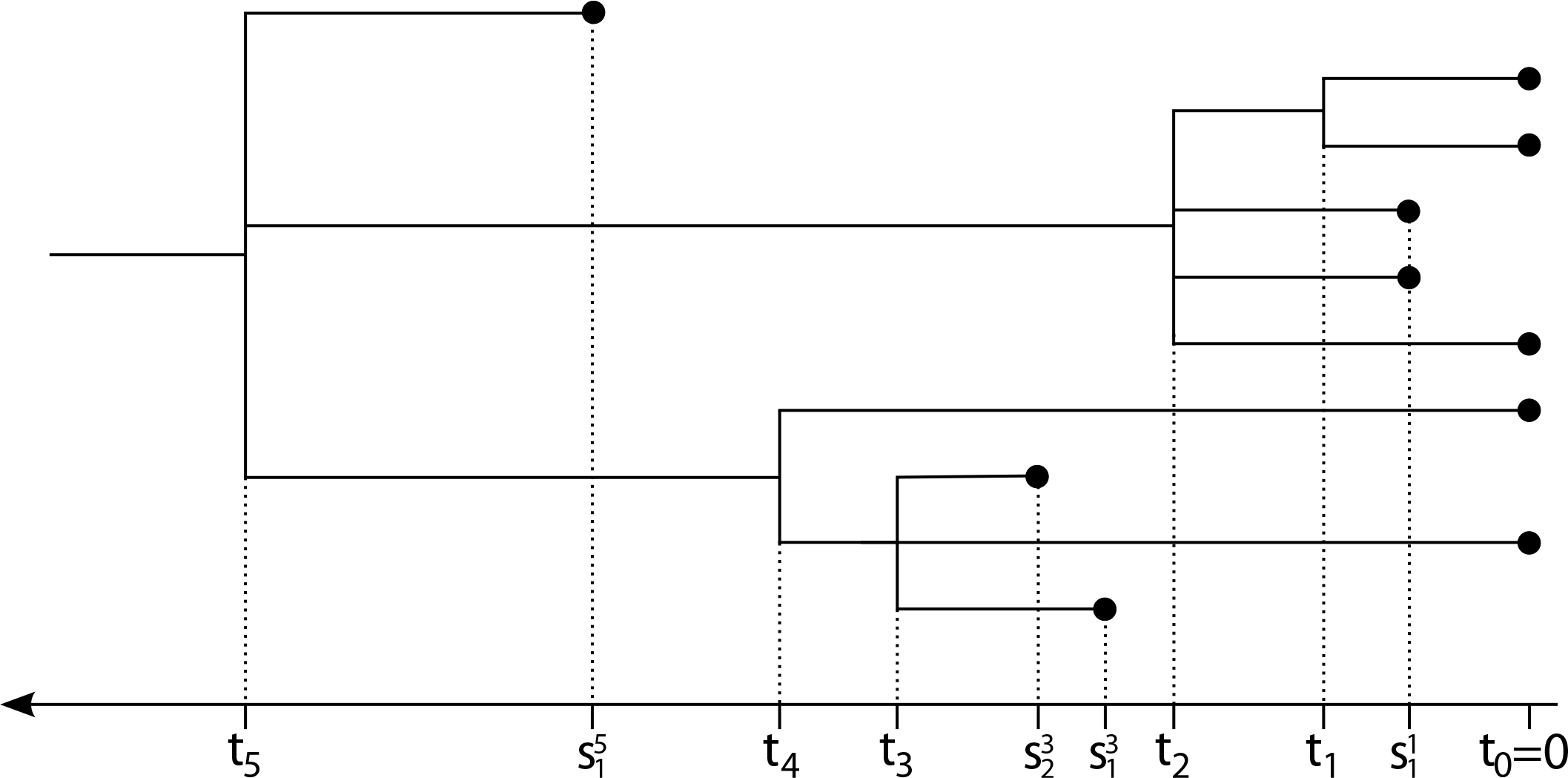
A serially-sampled multifurcating tree, with 10 tips in total. There are 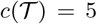 coalescent events. During the inter-coalescent interval [*t*_2_; *t*_3_), the number of extant lineages is (going back in time) *n*_2_ = *n*(1) = 3; *n*(2) = 4; and *n*(3) = 5. Using the composite-interval notation, assuming for example that *t*_4_ – *t*_3_ < ϵ < *t*_2_ – *t*_1_, the ensuing interval partition would have a 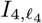 component, with *l*_4_ = 2 corresponding to the two intervals [*t*_3_; *t*_4_) and [*t*_4_; *t*_5_) having been joined together. There would be 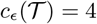 intervals in the merged interval partition

#### Inference using the multifurcating skyline plot

We evaluated the ability of the multifurcating skyline plot to estimate *N*_*e*_ by applying it to trees simulated under the Beta-coalescent process. We considered two different demographic histories (constant population size and exponential growth) and two values of the interval length threshold parameter ϵ. We used the same a parameter in both cases, namely *α* = 1.5. The results are shown in Fig. 5.

**FIG. 5.**
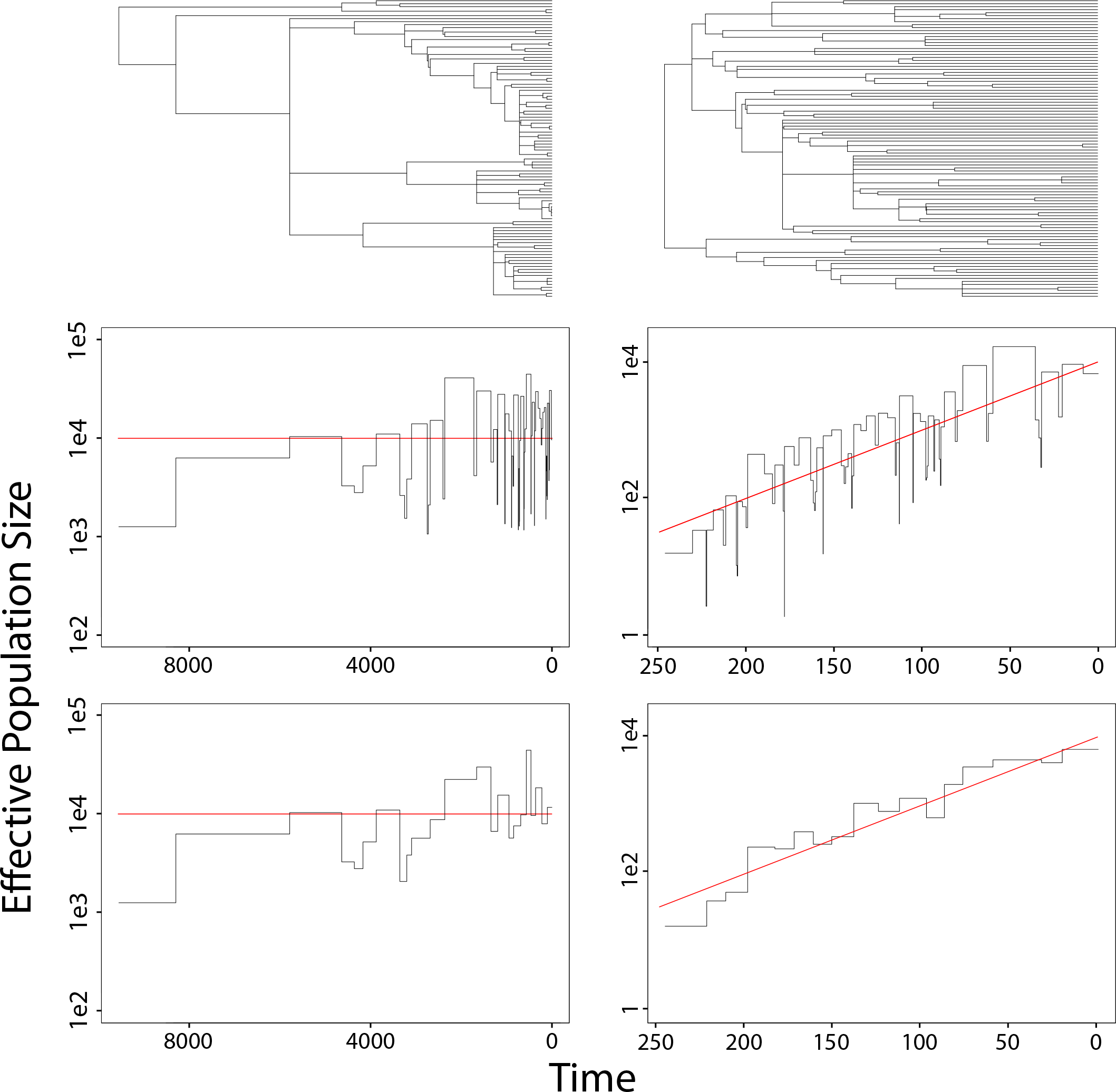
Top: Individual trees simulated with 100 tips under a Beta(0:5; 1:5)-coalescent, under constant (left) and exponential population size (right). Red lines represent the true effective population size function. Middle row: multifurcating skyline plots of the trees, with fixed *α* = 1:5 and with ϵ = 0. Bottom: Generalized multifurcating skyline plots, with threshold length parameter ϵ = 100 (left) and ϵ = 10 (right). Time is expressed in number of generations, and the estimated effective population size is in population size units.

The results are in line with well-known properties of skyline estimators (Drummond *et al.*, 2005; Strimmer and Pybus, 2001): they recover fluctuations in effective population size reasonably well, at the cost of being quite noisy. As expected, noise decreases as the interval merging parameter ϵ increases (Fig. 5.

#### Estimation of variance in offspring distributions

To illustrate potential uses of the multifurcating skyline plot, we apply the method to a serially-sampled molecular clock phylogeny estimated from Ebola virus genomes that were sampled during the epidemic in West Africa during 2015-2015. Specifically, we use the maximum clade credibility (MCC) tree resulting from a Bayesian phylogenetic analysis of 1610 Ebola virus sequences sampled between March 2014 and October 2015 in Sierra Leone, Liberia and Guinea. This MCC tree represents the output of the phylogeographic analysis published in (Dudas *et al.*, 2017) and is available from https://github.com/ebov/space-time. We also obtained weekly counts of Ebola cases during the epidemic from http://who.int/csr/disease/ebola/en/. These case count are used below as a proxy for the census number of infections, *N*_count_, over time.

The Ebola virus MCC phylogeny is strictly bifurcating. Since we do not attempt here to estimate multifurcating trees from gene sequence data, we used a heuristic approach to transform the binary tree into a non-binary one. Specifically, we aggregated binary nodes separated by short branch lengths into a multifurcation, as follows: for a given aggregation parameter *η* ≥ 0, whenever two internal nodes *x* and *y* existed at timepoints distant by less than *η* days, with *y* being a child of *x*, we collapsed *x* and *y* into a single node containing all lineages involved, located at the time of *x*. This procedure is performed by starting at the root node and proceeding to the most recent tip of the tree. For increasing values of *η*, more and more nodes will be aggregated, resulting in polytomies involving more and more lineages. Figure 6 shows the original binary phylogeny (*η* = 0) and two trees obtained using two different values of the aggregation parameter (*η* = 1 and *η* = 7.15 days). These “pseudo-multifurcating” trees are intended to illustrate the type of trees that might be expected if the true nature of epidemic transmission was known with certainty. Estimates of effective population size obtained by the skyline estimator were divided by the serial interval τ = 14.3 days, representing generation time, to obtain an estimate in units of population size.

**FIG. 6.**
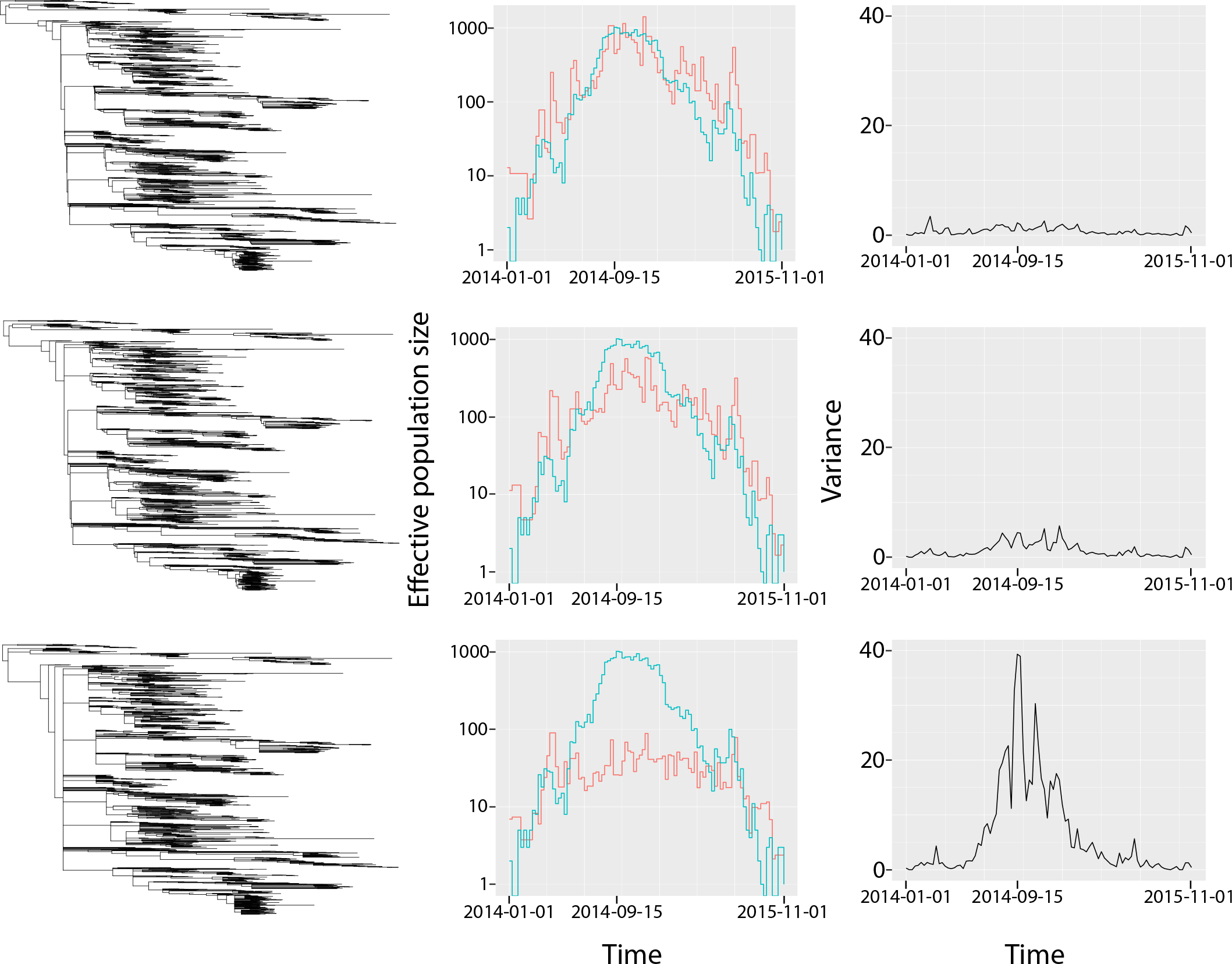
An application of the multifurcating skyline plot to a molecular clock phylogeny of the West Africa Ebola virus outbreak. The figure shows a visual comparison of phylogenies (left column), effective population size estimates (red line shown in middle column, population size units) and estimates of reproductive variance through time (right column) under three different values of *η*, the aggregation parameter. The top row shows *η* = 0 days, the middle row shows *η* = 1, and the bottom row shows *η* = 7:15 days. All graphs are plotted on the same timescale and the date of the epidemic peak (mid September 2014) is noted. The blue line in the middle column shows the number of reported Ebola virus cases during the outbreak, which is an estimate of *N*_count_

Importantly, varying the amount of multifurcation in the trees leads to different estimated *α* values and hence different skyline estimates of effective population size (Fig. 6). If no nodes are aggregated (i.e. *η* = 0) then the effective population size plot, as expected, agrees with previous estimates obtained using Kingman coalescent approaches (eg. (Dellicour et *al.*, 2018)) and is mostly proportional to *N*_count_, the weekly number of observed Ebola virus cases (Fig. 6 middle column). Using *N*_count_ (Fig. 6 blue lines) we are also able to estimate the variance in the reproductive number of infections over time, as σ^2^(*t*) = *N*_count_(*t*)/*N*_*e*_(*t*) (Fig. 6 right column). When *η* = 0 this variance is low and shows little change through time.

Next, we chose an aggregation parameter of *η* = 7.15 days (Fig. 6, bottom row), which is approximately half the estimated serial interval of Ebola virus during the epidemic (Wong et *al.*, 2017). This results in a tree with 744 internal nodes (compared to 1609 for the original binary tree). From this tree we estimated the maximum-likelihood value of *α* to be *α* = 1.29. This value of α is suggestive of high levels of heterogeneity in reproductive number, as indicated by the presence of several large polytomies in the aggregated tree. We then computed the corresponding multifurcating-skyline estimate of *N*_*e*_, with *α* = 1.29. The multifurcating skyline plot obtained when *η* = 7.15 (Fig. 6 bottom row) suggests that *N*_*e*_(*t*) is low and varies little throughout the epidemic. However, estimates of the variance in offspring number through time σ^2^(*t*) varies significantly through time, over orders of magnitude. The highest variance is observed at *t* = 38 weeks (approximately mid-September 2014), coinciding with the maximum value of *N*_count_. Thus, for values of α close to 1.0, the model interprets changes in the coalescence rate in the tree as variation in offspring variance rather than change in census population size. This is reflected in the power-law relationship we that we observe between reproductive variance and population size: σ^2^(*N*_count_) ∝: *N*^0.676^ (*R*^2^ = 0.92). This relationship is highly consistent with the assumption σ^2^(*N*) ∝ *N*^2-*α*^ that underlies the Beta(2*α*, *α*)-coalescent.

Lastly, we repeated the same procedure with a intermediate value of the aggregation parameter, *η* =1 day, which generated a more fully resolved tree with 1386 internal nodes. When *η* = 1 day, estimated effective population size varies more significantly over time of the epidemic than when *η* = 7.15 days, and estimates of reproductive variance through time are correspondingly lower (Fig. 6 middle row).

## III. DISCUSSION

We have demonstrated how σ-coalescents can be used to infer demographic trends from multifurcating trees, extending the range of “skyline plot”-like estimators beyond binary trees and the standard Kingman coalescent. Using simulated data, we showed that the a parameter of the `Beta` (2 – *α*, *α*) family of σ-coalescents can be accurately estimated, which interpolates between the Kingman coalescent (*α* = 2) and the Bolthausen-Sznitman coalescent (*α* = 1). We introduce the multifurcating skyline plot, which estimates effective population size from time-scaled non-binary trees, the tips of which may be sampled either longitudinally or concurrently. We validated this estimation method on simulated trees.

We applied the multifurcating skyline plot to a time-scaled tree estimated from Ebola virus genomes sampled during the 2014-2015 epidemic in West Africa. This analysis highlighted the importance of correctly estimating parameters of the σ distribution (specifically, in our case, the α parameter of the Beta(2 − *α*, *α*)-coalescent process), since these can have a major influence on the multifurcating skyline plot. Because we had only a binary tree at our disposal, we resorted to heuristic node-aggregation in order to generate a “pseudo-multifurcating” tree illustrating the kind of non-binary tree that might be generated by general Λ-coalescent processes. This is an unsatisfying solution and further work is required to implement the inference of multifurcating trees in Bayesian phylogenetic frameworks such as BEAST. This will necessitate the definition of tree operators capable of exploring the larger space of non-binary trees; whether this can be done without affecting computational performance remains an open question. Joint evaluation of molecular clock phylogenetic likelihoods and multifurcating tree prior probabilities has the potential to discriminate between genuine multifurcations, and short tree branches on which no mutations are observed.

Several popular Bayesian implementations of the skyline plot approach exist, which treat the skyline plot likelihood as a “prior distribution” on trees. These methods include the “Bayesian skyline plot” (Drummond *et al.*, 2005)], which uses the composite-interval procedure described in this paper to reduce noise, and the skyride (Minin *et al.*, 2008) and skygrid (Gill *et al.*, 2013) plots, which use sophisticated, time-aware, smoothing procedures to penalize population size changes. We expect similar approaches could be applied to the multifurcating-skyline plot. Recent work (Möller *et al.*, 2018) has illustrated that mis-specification of the tree prior during Bayesian phylogenetic inference may lead to erroneous estimation of other parameters, highlighting the need to implement a wider range of computationally-tractable tree priors. The multifurcating skyline plot may help to address this deficit. We hope it will prove useful in the analysis of populations that exhibit patterns of propagation leading to true polytomies in their phylogeny, such as superspreading (Lloyd-Smith *et al.*, 2005), strong selective regimes (Neher and Hallatschek, 2013) or even adaptive radiation (Schluter, 2000). The estimated parameters of the A distribution might be informative about these propagation processes, which are not taken into account in current phylodynamic approaches.

Although skyline plot methods based on the Kingman coalescent are commonplace in phylodynamic analysis, the actual values of estimated effective population size are not often interpreted directly. In some cases, “true”, census population size and effective population size are related only through nonlinear relations. For example, it was shown (Frost and Volz, 2010; Volz, 2012) that under SIR-type epidemiological models, effective population size through time *N*_*e*_(*t*) evolves proportionally to 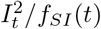, where *f*_*SI*_(*t*) is the rate at which susceptible individuals become infected (typically, *f*_*SI*_(*t*) ∝ *S*_*t*_*I*_*t*_/*N*). In this case, the relationship between effective population size *N*_*e*_(*t*) and disease incidence *I*_*t*_ at time *t* is different at the beginning and end of the epidemic. The multifurcating skyline methods will pose similar problems for interpretation: the actual value of the estimate is affected by the Λ distribution in a nonlinear (and not completely understood) way. Finally, it remains to be seen how specific Λ measures are related to actual characteristics of the population process generating the phylogeny. Existing results on Λ-coalescents ((Eldon and Wakeley, 2006; Schweinsberg, 2003)) give some indications of how specific Λ measures appear as scaling limits of discrete processes. It should also be noted that there are some indications ((Eldon et *al.*, 2015)) that changes in effective population size (for instance, exponential growth) can act as a confounding factor, albeit one that can be dealt with using multilocus data ((Koskela, 2018)).

## Acknowledgments

This work was supported by the European Research Council under the European Commission Seventh Framework Programme (FP7/2007-2013)/European Research Council grant agreement 614725-PATHPHYLODYN. Patrick Hoscheit has received the support of the EU in the framework of the Marie-Curie FP7 COFUND People Programme, through the award of an AgreenSkills fellowship under grant agreement n^○^ 267196.

The authors wish to thank Tim Vaughan for useful conversations that led to the overall improvement of this manuscript.

